# BTK inhibition enhances immunovirotherapy in glioblastoma via tertiary lymphoid structure modulation

**DOI:** 10.64898/2026.06.01.729275

**Authors:** Konstantina Kyritsi, Dong Zhu, Haocheng Ding, Gregory K. Friedman, Dongwen Lv, Meng Wang, Nahid F. Mivechi, Ravindra Kollhe, Theodore S. Johnson, Balveen Kaur, David H. Munn, Bangxing Hong

## Abstract

Glioblastoma (GBM) is a highly aggressive type of glioma that is resistant to immunotherapy and is associated with poor prognosis, largely due to its immunosuppressive tumor microenvironment. Bruton’s tyrosine kinase (BTK) is a non-receptor kinase that not only plays an important role in oncogenic signaling, particularly in tumor growth, but also regulates the activity of tumor-infiltrating myeloid cells, including dendritic cells, macrophages, and microglia in brain tumors. High BTK expression is associated with poor survival in patients with glioma. Oncolytic herpes simplex virus type 1 (oHSV)-derived virotherapy, a novel treatment strategy, has demonstrated effectiveness against GBM; however, its efficacy is limited by the tumor microenvironment. In this study, we found that BTK is predominantly expressed in GBM-infiltrating myeloid cells. Intratumoral injection of oHSV not only promotes infiltration of myeloid cells and T cells but also activates BTK in these myeloid cells, thereby limiting oHSV infection and replication in tumor cells. Combination treatment with BTK inhibitor ibrutinib improves anti-tumor efficacy of oHSV in both human GBM12 xenograft and syngeneic murine GSC005 models. Mechanistically, BTK inhibition increases oHSV-mediated tumor cell death (cleaved caspase-3) and cytotoxic CD8⁺ T cell infiltration, while decreasing tumor cell proliferation (Ki-67). BTK inhibition not only suppresses oHSV clearance by tumor-infiltrating microglia and macrophages but also reduces their pro-invasive effects on tumor cells. Addition of IDO inhibitor, an immune modulator, further prolongs survival in tumor-bearing mice in a syngeneic GBM model. Single-cell mRNA sequencing (scRNA-seq) analysis indicates that combination treatment modifies key signaling pathways in both tumor-infiltrating myeloid cells (macrophages and microglia) and CD8⁺ T cells. Further analysis shows that BTK inhibition, with or without IDO inhibition, promotes the formation of tumor-infiltrating tertiary lymphoid structures (TLS) during intratumoral oHSV treatment, subsequently remodeling T cell, NKT cell, and monocyte–macrophage populations. These results indicate that BTK inhibition exerts multifaceted effects in enhancing the anti-tumor efficacy of oHSV therapy.

## Introduction

Glioblastoma (GBM) is the most common primary malignant brain tumor, and outcomes are poor with a median survival of approximately 15 months from diagnosis^1^. Resistance to standard therapies, including chemotherapy, and radiotherapy, develops frequently, highlighting a critical unmet need for novel therapeutic strategies that can overcome treatment resistance.

Bruton’s tyrosine kinase (BTK), a member of the BTK family of non-receptor tyrosine kinases, plays an important role in multiple cellular processes, including angiogenesis, proliferation, inflammation, and apoptosis^2–5^. BTK is highly expressed in several cancer types, including prostate cancer, B-cell malignancies, and leukemia^6^. Ibrutinib, a selective BTK inhibitor, has been approved by the U.S. Food and Drug Administration (FDA) for the treatment of multiple malignancies, including chronic lymphocytic leukemia (CLL), mantle cell lymphoma (MCL), diffuse large B-cell lymphoma, follicular lymphoma, and multiple myeloma^7^. Therapeutically, ibrutinib has been shown to improve fibrotic responses, promote T-cell restoration, and induce tumor regression^8,9^. In GBM, ibrutinib has been reported to inhibit tumor growth by regulating BMX–STAT3 signaling in glioma stem cells, leading to reduced tumor cell proliferation, migration, invasion, and autophagy^10^.

In addition to its well-established oncogenic role in hematological malignancies, BTK has recently been recognized as an important regulator of immune function in solid tumors, particularly through its activity in tumor-infiltrating myeloid cells, including antigen-presenting cells^11^. Oncolytic virotherapy has emerged as a novel immunotherapeutic approach capable of enhancing immune cell infiltration in immunologically “cold” tumors^12^. Herpes simplex virus–based oncolytic virotherapy (oHSV) has achieved clinical validation, exemplified by the approval of talimogene laherparepvec (T-VEC) for the treatment of unresectable melanoma^13–15^. In parallel, multiple next-generation oHSV platforms are currently under clinical investigation for a range of solid malignancies, including GBM^16,17^. In this study, we investigate the role of brain myeloid BTK signaling in modulating the therapeutic efficacy of intratumoral oHSV therapy in GBM.

## Materials and Methods

### Cell lines, oncolytic virus and reagents

Patient-derived GBM12 and GBM28 cells, are cultured in Dulbecco’s Modified Eagle Medium (DMEM) added 2% fetal bovine serum (FBS), and 1% Penicillin/Streptomycin. Mouse GSC005 were cultured as spheres in DMEM/F12 media (Sigma-Aldrich D80620) supplemented with 2 mM L-glutamine, 1% N2 supplement, 0.5% penicillin streptomycin, recombinant human epidermal growth factor (EGF) (20 ng/mL), recombinant human FGF-basic (20 ng/mL) and puromycin. Accutase was used for spheres dissociation and passaging. BV2 murine microglia cell line was kept in culture with DMEM supplemented by 10% FBS and 1% Penicillin/Streptomycin. All cells were incubated at 37^0^C, validated to lack of contamination and maintained below thirty passages before the experiment started. All cells tested and confirmed negative for mycoplasma (Mycoplasma PCR Detection Kit, #G238, Applied Biological Materials, BC, Canada). Dr. Balveen Kaur (Louisiana State University) kindly provided GBM cells. Indoximod (1-methyl-D-tryptophan, clinical grade) was obtained from Lumos Pharma (New Link Genetics) and Ibrutinib (cat. # HY-10997) was purchased from Mechem Express. oHSV used in this study is PKR-shRNA virus (oHSV-shPKR and oHSV-mshPKR) generated in the lab^18^, which contains double mutation of ICP6 and gamma 34.5 genes, and expresses PKR-shRNA to increase virus infection and replication capacity in tumor cells.

### Infection and tumor cell viability *in vitro*

The infectivity of oHSV were performed in human GBM cells with different multiplicities of infection (MOI) (0.001–0.5), to determine the best MOI to infect the cells (MOI=0.05) for the combination with BTK inhibitor. Virus-infected GBM cells were quantified as GFP+ tumor cells by the Incucyte SX5 Live-Cell Analysis System from Sartorius. Tumor cells viability was analyzed and quantified by Aqua live/dead^TM^ Fixable Near IR (780) viability kit (Invitrogen, L34994) and data were analyzed using a flow cytometer (NovoCyte Quanteon, Santa Clara, CA).

### *q*PCR

RNA was isolated from GSC005 tumors using *q*PCR Qiagen RNeasy Mini Kit (74104, Qiagen, Germantown, MD) following the manufacturer’s protocol. To synthesize cDNA, 2 mg of quantified RNA was reverse transcribed using the High-Capacity cDNA Reverse Transcription Kit (Applied Biosystems, Foster City, CA) using random hexamers. Primers were designed via Primer3 software and synthesized from Integrated DNA Technologies, Inc. (IDT, Coralville, IA). *q*PCR was performed using qPCRBIO SyGreen Mix Lo-ROX (Genesee Scientific, San Diego, CA) in QuantStudio 3 Real-Time PCR System (Thermo Fisher Scientific, Waltham, MA) in 96-well plates with cycling conditions as such: initial denaturation at 95^0^C for 2 mins; amplification cycle: denaturation at 95^0^C for 15 s, annealing at 60^0^C for 20 s, and extension at 72^0^C for 5 s. *GAPDH* were used as housekeeping controls and 2(DDCt) method was used to determine differential fold change in expression.

### Animal models

Animal experiments were performed consonantly with protocols approved by Augusta University (protocol #: 2023-1098). C57/BL6 (JAX:000664) and nude mice (JAX: 002019) were purchased from the Jackson Laboratory, Bar Harbor, ME. All mice, five per cage, were housed in room-controlled temperature on a 12:12-h light cycle with access to food, standard chow pellet diet, and water. Before the mice underwent stereotactic intracranial injection, anesthesia was performed with intraperitoneal injection of 0.2 mL of stock solution containing the following drugs: ketamine HCl (25 mg/mL) and xylazine (2.5 mg/mL) diluted in distilled water. Mice were fixed in a stereotactic device (David Kopf Instruments, Tujunga, CA), the surgical site anointed with 70% ethyl alcohol and a skin incision was performed over the midline. Over the right hemisphere at a location 2 mm lateral and 1 mm anterior to bregma, GBM12, GSC005 cells or oHSV in 2 μL PBS were intra-cranially injected using needles (Hamilton 80300 for cell implantation and Hamilton 80000 for virus) at a depth of 3.5 mm and at a rate of 0.4 μL/min using autoinjectors (KD Scientific Inc, Holliston, MA). Slowly, needles were removed, and skin was sutured using a 5-0 nylon thread. Seven days after tumor cells injection, mice were treated intra-tumorally with sterile PBS or 2 × 10^5^ PFU of oHSV and started daily treatment by intraperitoneal injection with Ibrutinib (16mg/kg/day), which was continued for 14 days. Animals were observed and euthanized at the indicated time points and brains were harvested.

### Immunofluorescence and immunohistrochemistry staining

Human GBM paraffin tissue array was purchased from TissueArray.Com LLC (GL722b). Mouse brain tumors were collected to prepare paraffin blocks and sections. After deparaffinization and antigen retrieval, the sections were permeabilized with 0.04% triton-X and blocked with 2% goat serum and incubated with primary antibodies (BTK, CD68, CD11c, CD8, Foxp3, cleaved caspase-3, Ki-67, CD3 and MAdCAM-1) overnight at 4^0^C. The day after, the sections were washed and incubated with secondary antibodies for 1 hour. Slides were fixed with cover slips using Fluor mount-G^TM^, with DAPI (Invitrogen, E142914) solution, images were taken by fluorescence microscopy (Nikon Eclipse Ts2) and analyzed using ImageJ software.

### Flow cytometry

For cell surface staining, cells were washed with PBS and blocked with an Fc blocker (BD Biosciences, San Jose, CA). Fluorochrome-labeled antibodies (CD45, CD11b, P2Y12, CD163, CD8, and BTK) were obtained from BD Biosciences (Franklin Lakes, NJ), added, and stained for 30 minutes as previously described^18^. All samples were analyzed using a CytoFlex flow cytometer (Beckman Coulter, CA).

### Single-Cell mRNA sequencing

CD45+ cells were isolated from tumors treated with, PBS, oHSV and ibrutinib, or oHSV and ibrutinib and indoximod. mRNA libraries were constructed using the 10x Genomics Chromium Next GEM Single Cell 5’ HT Reagent Kits v2 (Dual Index). Raw sequencing data files were demultiplexed into FASTQ files and analyzed using the Cell Ranger algorithm (10x Genomics). The filtered count matrices and filtered contig V(D)J annotations were analyzed with R (v 4.2) using Seurat and Bioconductor packages. The low-quality cells were filtered out retaining cells with detected gene numbers >200, and mitochondrial genes <15%. Genes that were expressed by less than 3 cells were rejected.

### Statistical analysis

All quantitative results are presented as means ± standard deviation (SD). Statistical differences between two groups were assessed using the Mann-Whitney U test or Student’s *t*-test. For comparisons involving more than two groups, ANOVA was employed. Statistical analyses were conducted using Prism 5 software (GraphPad Software, Inc., La Jolla, CA). A p-value of less than 0.05 was considered statistically significant.

## Results

### BTK is predominantly expressed in GBM-Infiltrating myeloid cells

BTK is expressed not only in tumor cells^19^ but also in tumor-infiltrating myeloid cells^20^. To evaluate the potential of targeting BTK signaling in GBM, we first examined BTK expression in tumor samples from GBM patients. Given that the majority of immune cells infiltrating GBM are of myeloid origin^21^—including macrophages, microglia, and dendritic cells—we performed co-immunostaining of BTK with CD68 (a marker for monocytes, macrophages, and microglia), CD11c (a dendritic cell marker), and T cell markers (CD8 and Foxp3) on tissue sections from GBM tumors and normal brain tissue (**Fig. 1a–b**).

**Fig. 1.**
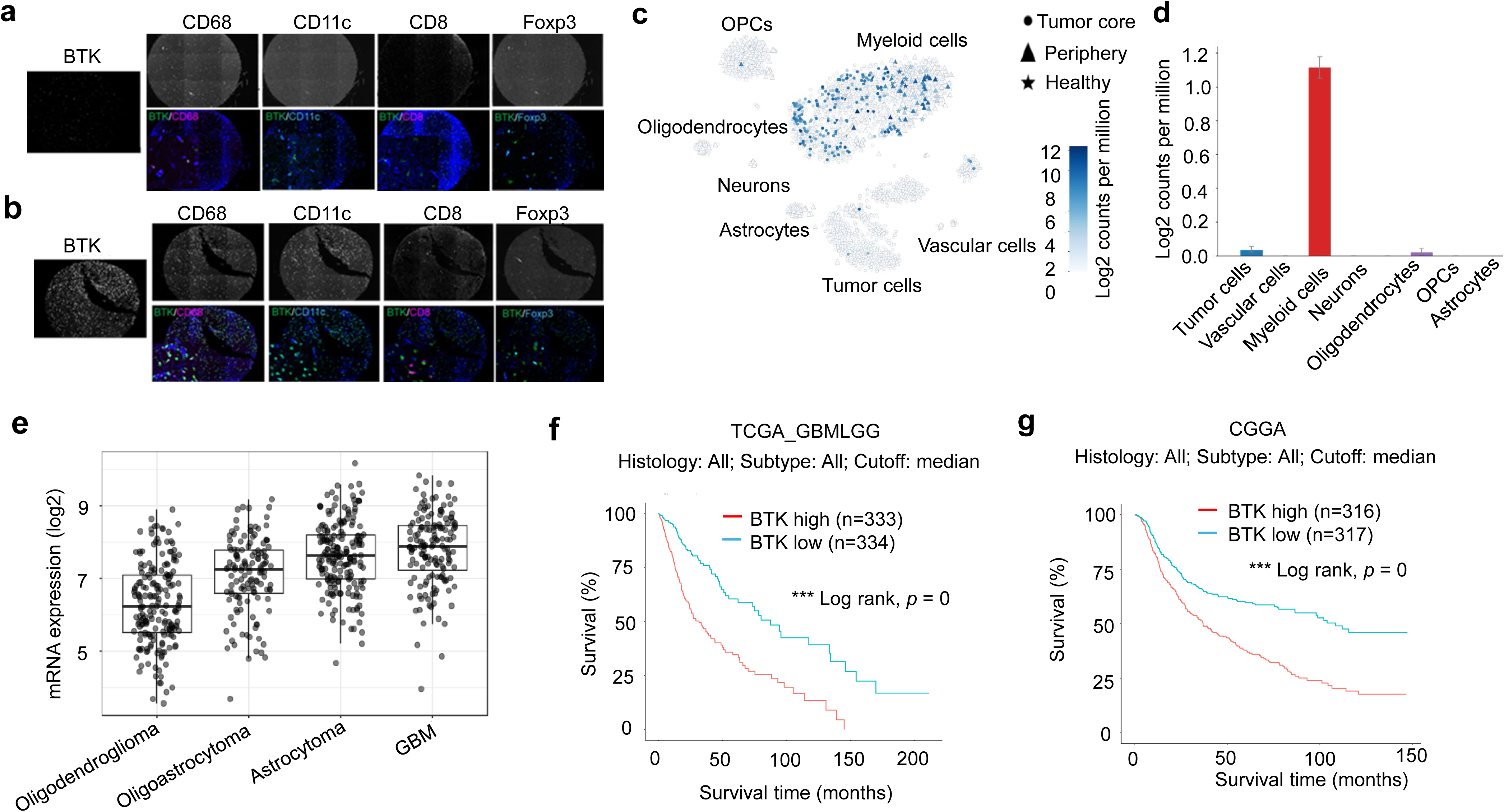
BTK expression in human glioblastoma and mouse models. (**a, b**) Representative multicolor immunofluorescence staining of BTK, CD68, CD11c, CD8, and Foxp3 in human normal brain (**a**) and glioblastoma (GBM) specimens (**b**). Images demonstrate that BTK expression is predominantly localized within CD68⁺ and CD11c⁺ myeloid populations, with minimal expression in CD8⁺ or Foxp3⁺ lymphocytes. (**c, d**) scRNA-seq analysis of GBM patient tumors. UMAP visualization (**c**) identifies major tumor and immune cell clusters, and corresponding bar plots (**d**) quantify BTK expression levels across annotated cell types, confirming enrichment in myeloid compartments. (**e**) Comparative analysis of BTK expression across GBM and other brain tumor types, demonstrating elevated BTK expression in GBM relative to lower-grade gliomas and non-glioma brain tumors. (**f, g**) Kaplan–Meier survival analyses showing that high BTK expression is significantly associated with reduced overall survival in GBM patients from The Cancer Genome Atlas (TCGA, **f**) and the Chinese Glioma Genome Atlas (CGGA, **g**) cohorts.

Our results demonstrated that BTK staining was seen in GBM tissue but not in normal brain tissue (**Fig. 1a–b**). Notably, tumor-infiltrating CD68⁺ and CD11c⁺ cells exhibited higher levels of BTK expression than CD8⁺ T cells and Foxp3⁺ regulatory T (Treg) cells (**Fig. 1b**), indicating that BTK is preferentially expressed in myeloid cell populations within the GBM microenvironment.

To further investigate the role of BTK in human GBM, we analyzed publicly available single-cell RNA sequencing (scRNA-seq) datasets (**Fig. 1c–e**). The analysis revealed that BTK expression is predominantly enriched in CD45⁺ immune cells within the tumor microenvironment, particularly among myeloid populations, including macrophages, monocytes, dendritic cells, and neutrophils (**Fig. 1c–d**). In contrast, tumor cells exhibited relatively low levels of BTK expression, consistent with previous reports indicating that BTK is restricted to a small subset of glioma stem-like cells^10^.

Comparative analysis further showed that GBM tumors exhibit higher overall BTK expression than other subtypes, including oligodendroglioma, oligoastrocytoma, and astrocytoma (**Fig. 1e**). Importantly, elevated BTK expression was positively correlated with poor prognosis in GBM patients, as demonstrated in both the TCGA (**Fig. 1f**) and CGGA (**Fig. 1g**) datasets.

### Intratumoral oHSV treatment modulates BTK signaling in tumor-Infiltrating myeloid cells

Oncolytic herpes simplex virus (oHSV) has been developed as a therapeutic strategy for glioblastoma (GBM)^16,17^. To determine whether intratumoral oHSV administration modulates BTK signaling in tumor-infiltrating myeloid cells, we first assessed BTK expression in both human GBM12 and murine GSC005 orthotopic GBM models established in Nude and C57BL/6J mice, respectively. We observed substantial infiltration of BTK⁺ cells in both human and murine GBM tumors compared to adjacent brain tissue (**Fig. 2a**). Flow cytometry analysis of GSC005 tumors in C57BL/6J mice demonstrated that tumor cells (GSC005-GFP⁺) express low levels of BTK, whereas tumor-infiltrating microglia (P2Y12⁺) and macrophages (CD163⁺) exhibit high BTK expression (**Fig. 2b**), indicating that BTK is predominantly enriched in myeloid cell populations within the tumor microenvironment.

**Fig. 2.**
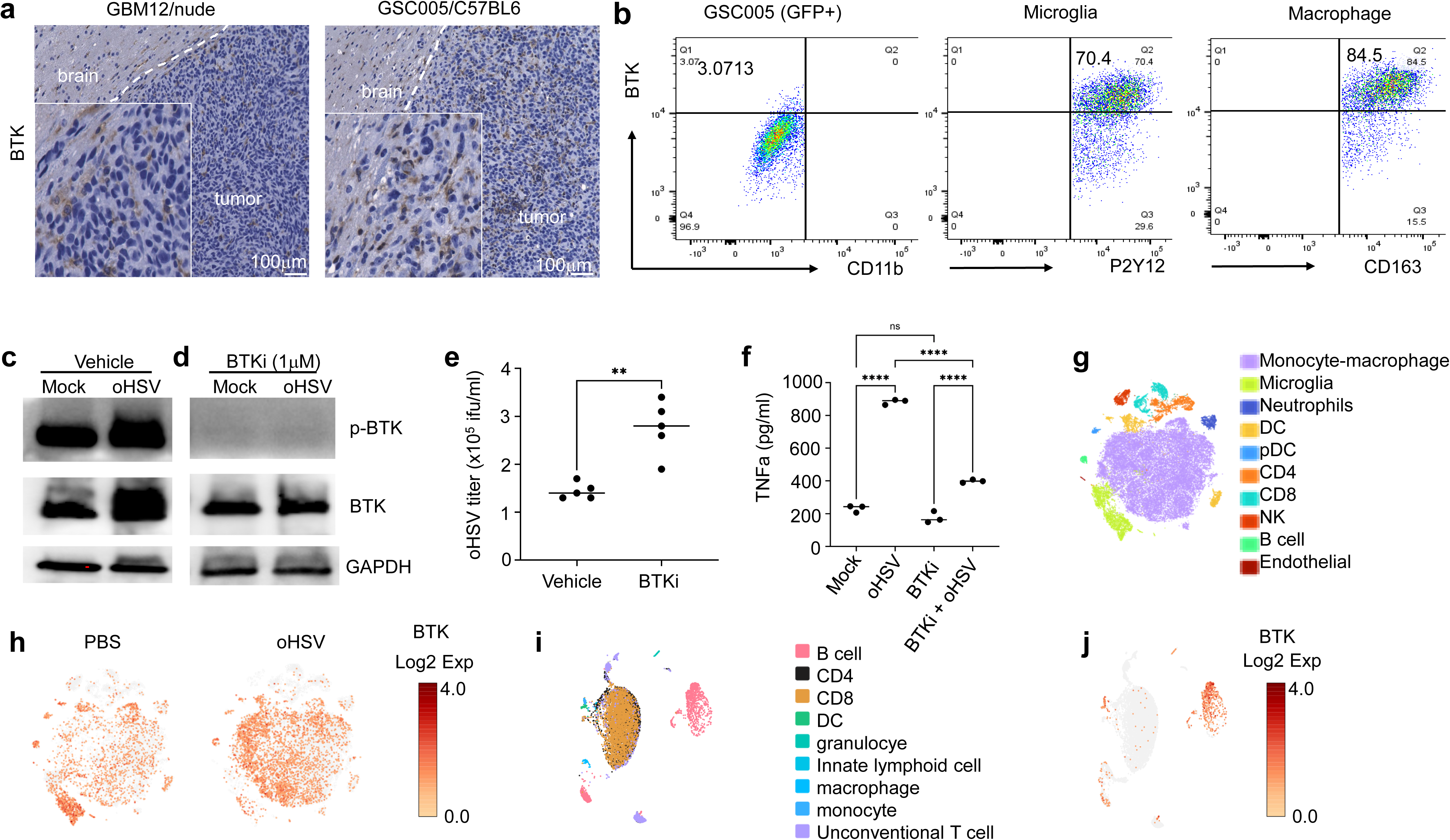
oHSV) treatment upregulates BTK expression in GBM tumor microenvironment. (**a**) IHC staining of BTK in orthotopic human GBM12 model in nude mice and murine GSC005 model in C57BL/6J mice. (**b**) Flow cytometry analysis of BTK expression in tumor cell (GFP+), tumor-infiltrated microglia (P2Y12) and macrophage (CD163) in GSC005 model. (**c-f**) Mock- or oHSV-infected GSC005 were cocultured with BV2 microglia cells and treated with vehicle or ibrutinib (BTKi). BTK activation was analyzed by western blot (**c, d**). oHSV replication after BTK inhibition was determined by titration assay (**e**). n=5, ** *p*<0.01. ns=no significance. TNFa secretion in the coculture was analyzed by ELISA (**f**). n=5, **** *p*<0.001. ns=no significance. (**g–j**) scRNA-seq analysis of CD45⁺ immune cells isolated from murine 005 tumors (**g, h**) and tumor-draining lymph nodes (TDLNs) (**i, j**). tSNE plots and corresponding quantifications reveal dynamic changes in BTK expression across immune subsets following oHSV treatment. (**i, j**) UMAP plots depicting BTK expression patterns in TDLNs after intratumoral oHSV administration, highlighting increased BTK expression in specific immune populations.

We next investigated whether oHSV treatment influences BTK signaling in these cells. Co-culture of oHSV-infected GSC005 cells with BV2 microglial cells resulted in a significant increase in BTK expression compared to co-culture with mock-infected GSC005 cells (**Fig. 2c**). Treatment with the BTK inhibitor ibrutinib effectively abolished BTK activation (phospho-BTK) in both GSC005 and oHSV-infected GSC005 co-culture systems (**Fig. 2d**). Given the known role of microglia and macrophages in clearing viral infections, we further evaluated the impact of BTK inhibition on oHSV clearance. Viral titration assays revealed that treatment with ibrutinib significantly reduced oHSV clearance (**Fig. 2e**) and decreased TNF-α secretion in the co-culture system (**Fig. 2f**), suggesting that BTK activity contributes to antiviral responses mediated by myeloid cells.

Finally, we assessed the *in vivo* effects of intratumoral oHSV administration in GSC005 tumor-bearing mice (**Fig. 2g–h**). Single-cell RNA sequencing (scRNA-seq) analysis demonstrated that intra-tumor injection of oHSV significantly upregulates BTK expression in tumor-infiltrating myeloid populations, including monocytes/macrophages and microglia (**Fig. 2h**). In contrast, analysis of tumor-draining lymph nodes revealed that BTK is primarily expressed in B cells, with lower expression observed in myeloid populations such as dendritic cells, macrophages, and monocytes (**Fig. 2i–j**).

### BTK inhibition suppresses GBM neurosphere growth and enhances oHSV-mediated tumor lysis

Previous studies have shown that a subset of glioblastoma (GBM) stem-like cells is responsive to growth inhibition by ibrutinib, a BTK inhibitor^10^. To determine the effective concentration of ibrutinib in GBM neurospheres, we first performed dose-response analyses using GBM12 and GBM28 neurosphere cultures (**Fig. 3a–b**). Cells were treated with increasing concentrations of ibrutinib (2.5, 5, 10, and 12.5 µg/mL), and tumor cell growth was monitored using live-cell imaging on the Incucyte system over 48–60 hours. The data indicated that 2.5 µg/mL of ibrutinib was sufficient to inhibit growth in both GBM12 and GBM28 neurospheres (**Fig. 3a–b**). Next, we evaluated the effect of combining ibrutinib with oHSV in these human GBM neurospheres. Co-treatment did not significantly affect oHSV infection efficiency (**Fig. 3c–d**), indicating that BTK inhibition does not interfere with viral entry or replication. To assess neurospheres growth and cell death, GBM12 and GBM28 neurospheres were analyzed by Incucyte live-cell imaging (growth) and flow cytometry (cell death) (**Fig. 3e–h**). The combination of ibrutinib and oHSV significantly inhibited neurosphere growth (**Fig. 3e, g**) and increased tumor cell death (**Fig. 3f, h**) in both GBM12 and GBM28 neurospheres. These results demonstrate that BTK inhibition not only suppresses GBM stem-like cell proliferation but also enhances the cytolytic activity of oncolytic virus therapy, supporting the potential therapeutic synergy of this combinatorial approach.

**Fig. 3.**
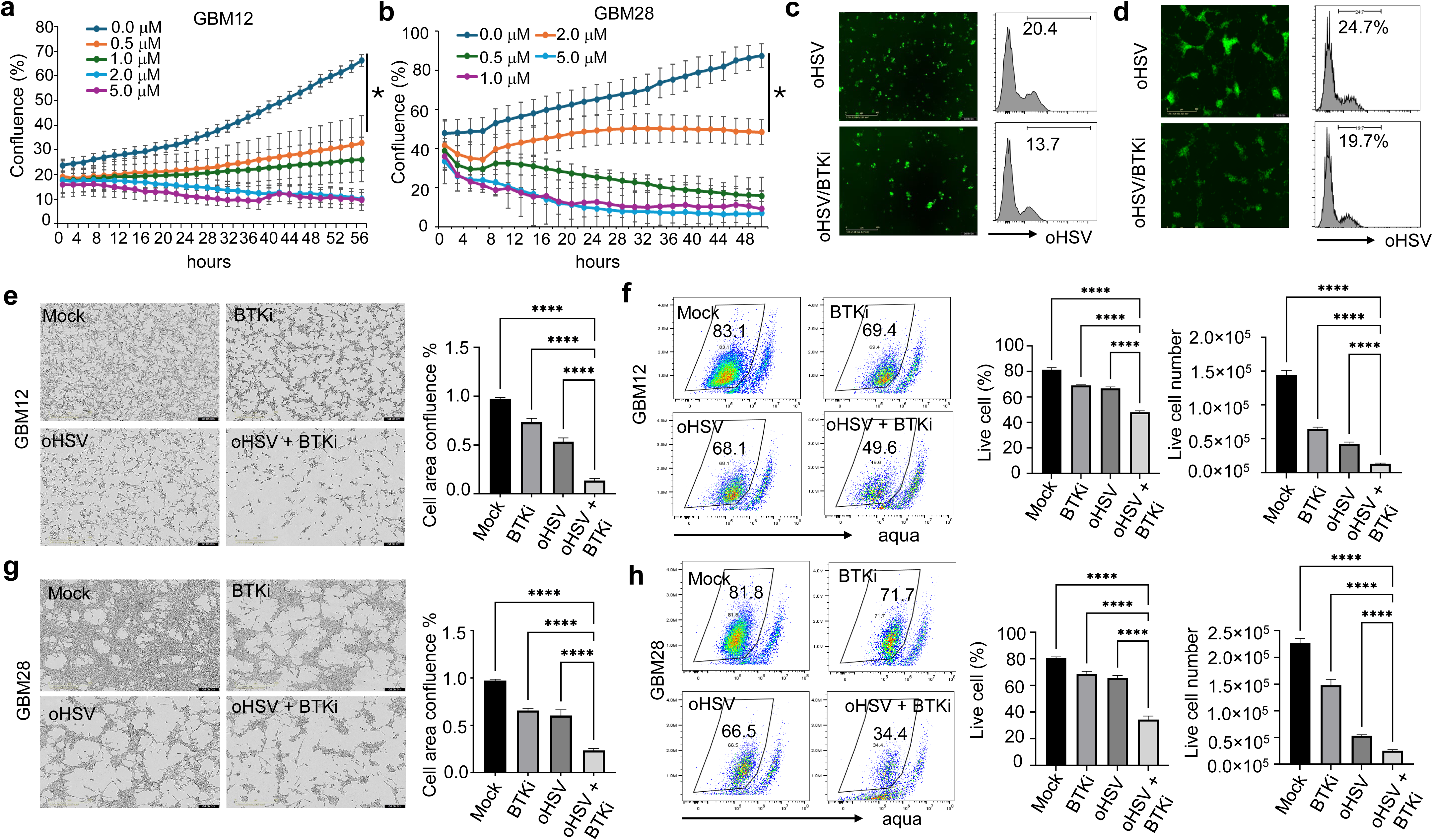
BTK inhibition enhances oHSV-mediated tumor cell killing *in vitro*. (**a, b**) Growth kinetics of GBM12 (**a**) and GBM28 (**b**) neurospheres treated with the BTK inhibitor ibrutinib, monitored using the Incucyte live-cell imaging system. (**c, d**) Flow cytometric analysis of oHSV(Multiplicity of infection, MOI=0.01) infection efficiency in GBM12 (**c**) and GBM28 (**d**) cells in the presence or absence of 0.5μM ibrutinib. (**e–h**) Quantification of tumor cell death following treatment with oHSV (MOI=0.01) ± 0.5μM ibrutinib in GBM12 (**e, f**) and GBM28 (**g, h**) models. Cell death was assessed by Incucyte-based cytotoxicity assays (**e, g**) and Aqua live/dead staining followed by flow cytometry (**f, h**), indicating significantly increased cytotoxicity with combination treatment. Data are presented as mean ± SD from triplicate experiments; * p < 0.05, ****p < 0.0001.

### Ibrutinib enhances the anti-Tumor activity of oHSV in an orthotopic human GBM model independent of adaptive immunity

Given that ibrutinib enhances oHSV-mediated tumor cell lysis *in vitro* in human GBM neurospheres, we next evaluated the therapeutic efficacy of the combination *in vivo* using an orthotopic human GBM model in nude mice, which lack functional T cells, allowing assessment independent of adaptive immunity. GBM12 cells were implanted intracranially in nude mice, followed by intratumoral administration of oHSV and systemic ibrutinib (i.p.) (**Fig. 4a**). The combination treatment significantly prolonged survival of tumor-bearing mice compared to either monotherapy (**Fig. 4b**). Immunofluorescence analysis of cleaved caspase-3 (CC3), an indicator of apoptosis, demonstrated that ibrutinib enhanced tumor cell death, while by immunohistochemistry staining of proliferation marker (Ki-67) revealed a reduction of the tumor growth (**Fig. 4c-d**).

**Fig. 4.**
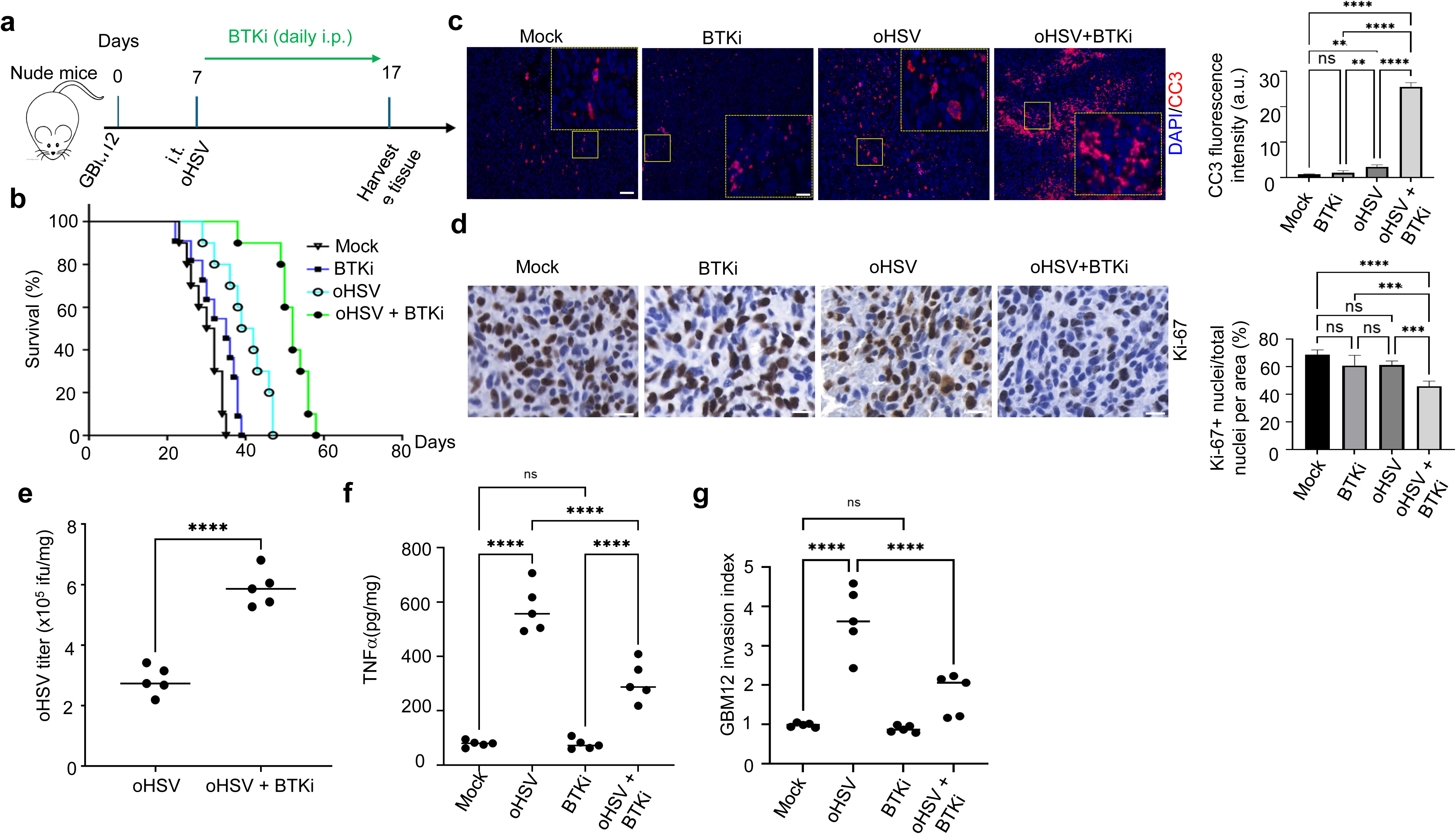
BTK inhibition enhances the therapeutic efficacy of oHSV in human GBM *in vivo*. **(a)** Schematic representation of the intracranial GBM12 xenograft model in immunodeficient (nude) mice and treatment regimen with oHSV and the BTK inhibitor ibrutinib. **(b)** Kaplan–Meier survival curves demonstrating prolonged survival in mice treated with the combination therapy compared to monotherapy or control groups. n=10, *p*=0.024 **(c)** Immunofluorescence staining of cleaved caspase-3 (CC3) in tumor sections, indicating enhanced apoptosis in the combination treatment group. (**d**) Immunohistochemistry staining of tumor cell proliferation marker (Ki-67) in tumor sections, indicating decrease tumor cell growth in the combination treatment group. (**e**) Virus titration of tumor after treatment with oHSV and BTKi ibrutinib. n=5, **** *p*<0.0001. (**f**) TNF-α secretion in the tumor lysate after treatment with oHSV and BTKi ibrutinib. n=5, **** *p*<0.0001. ns=no significance. (**g**) The effect of microglia (P2Y12+) isolated from the tumor on GBM12 cells invasion ex vivo was analyzed using transwell assay. n=5, **** *p*<0.0001. ns=no significance.

Although nude mice lack functional T cells, they retain functional myeloid populations within the tumor, including microglia and macrophages, which are known to efficiently clear oHSV infection. We observed that oHSV co-treatment with ibrutinib reduced viral clearance (**Fig. 4e**). Consistent with this, TNF-α secretion from tumors decreased following combination therapy (**Fig. 4f**). Tumor-infiltrating myeloid cells, including tumor-infiltrated microglia, contribute to GBM invasiveness^22^. P2Y12^+^ microglia isolated from combination-treated tumors exhibited a reduced ability to promote GBM12 neurosphere invasion *ex vivo* compared to those from oHSV-treatment group (**Fig. 4g**).

These results indicate that BTK inhibition enhances oHSV anti-tumor efficacy *in vivo* by modulating the function of tumor-infiltrating myeloid cells in oHSV clearance and tumor cell invasion, independent of adaptive immune responses.

### Combination of oHSV with ibrutinib, with or without IDO Inhibition, enhances tumor cell lysis and CD8⁺ T cell infiltration in the murine GSC005 model

To investigate whether ibrutinib enhances the anti-tumor efficacy of oHSV through activation of adaptive immunity, we employed the syngeneic murine GSC005 GBM model. Combination treatment with ibrutinib and oHSV significantly prolonged survival of tumor-bearing mice compared to either monotherapy (**Fig. 5a–b**). Immunofluorescence analysis of cleaved caspase-3 (CC3) revealed that ibrutinib markedly increased oHSV-mediated tumor cell lysis in vivo (**Fig. 5c**). Because tumor proliferation is a major driver of GBM progression, we assessed Ki-67 expression as a marker of tumor cell growth in intracranial GSC005 tumors. The combination of ibrutinib and oHSV significantly reduced Ki-67⁺ tumor cell populations and simultaneously enhanced CD8⁺ T cell infiltration compared to single treatments (**Fig. 5d and sFig.1**), demonstrating that BTK inhibition promotes anti-tumor immunity and suppresses tumor proliferation.

**Fig. 5.**
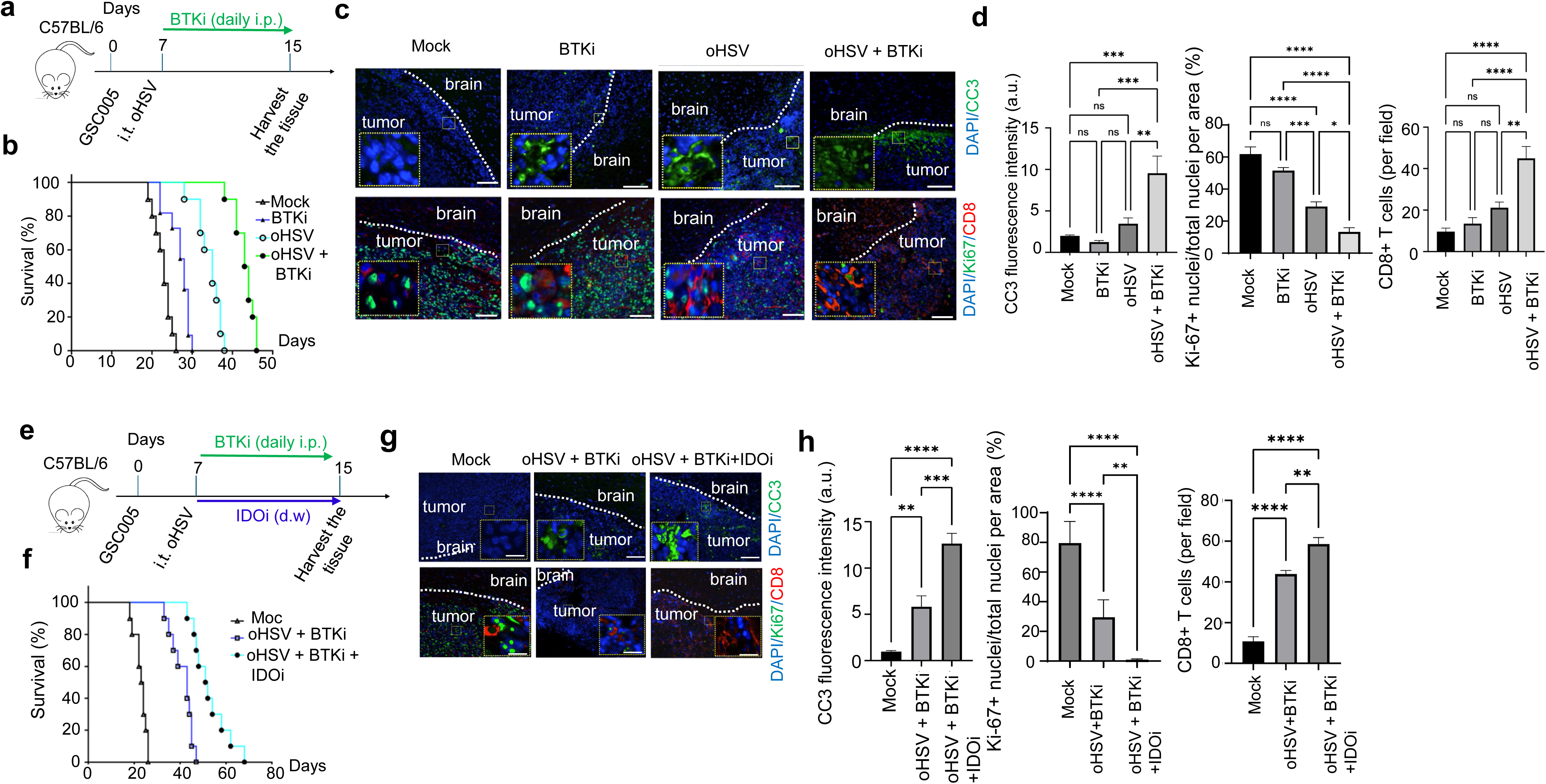
Combined BTK and IDO inhibition enhances oHSV-mediated anti-tumor responses in murine GSC005 GBM. (**a**) Experimental schematic of the syngeneic murine GSC005 GBM model treated with oHSV and ibrutinib. (**b**) Kaplan–Meier survival analysis showing improved survival with combination therapy. n=10, *p*=0.027. (**c, d**) Immunostaining of CC3 demonstrating increased tumor cell apoptosis following treatment. Immunofluorescence staining of Ki-67 (proliferation marker) and CD8 (cytotoxic T cells), indicating reduced tumor proliferation and increased T cell infiltration. n=5, * *p*<0.05, ** *p*<0.01, *** *p*<0.005, **** *p*<0.0001. ns=no significance. (**e**) Schematic of triple-combination therapy incorporating oHSV, ibrutinib, and the IDO inhibitor indoximod. (**f**) Kaplan–Meier survival analysis demonstrating further survival benefit with triple therapy. n=10, *p*=0.031. (**g, h**) CC3 staining showed enhanced apoptosis in tumors treated with triple combination therapy. Ki-67 and CD8 staining indicated further suppression of tumor proliferation and increased cytotoxic T cell infiltration. n=5, ** *p*<0.01, *** *p*<0.005, **** *p*<0.0001.

Indoleamine 2,3-dioxygenase (IDO) is an immunoregulatory enzyme frequently upregulated in tumors, including GBM, where it suppresses antigen presentation and dampens anti-tumor immunity^23^. Blockade of IDO signaling using indoximod enhances dendritic cell activation during oncolytic virotherapy^23^. Recent studies indicate that BTK regulates IDO signaling in antigen-presenting cells via the BTK–IDO–mTOR axis, and inhibition of this pathway boosts anti-tumor T cell immunity by promoting monocyte-to-dendritic cell differentiation^11^. We next evaluated whether addition of the IDO inhibitor indoximod could further enhance the efficacy of ibrutinib plus oHSV (**Fig. 5e**). Co-administration of indoximod further prolonged survival of tumor-bearing mice compared to the ibrutinib–oHSV combination alone (**Fig. 5f**). Immunofluorescence analysis of CC3 demonstrated increased tumor cell death with triple therapy (**Fig. 5g**). Furthermore, triple treatment significantly reduced tumor cell proliferation (Ki-67) and increased CD8⁺ T cell infiltration compared to dual therapy (**Fig. 5h**). These data indicate that IDO inhibition reinforces the anti-tumor and immunostimulatory effects of the ibrutinib–oHSV combination.

### Combination of oHSV with BTK and IDO inhibition promotes tertiary lymphoid structure formation in the GSC005 model

To investigate whether the enhanced CD8⁺ T cell response observed during BTK inhibition was associated with improved tumor antigen presentation, we focused on tertiary lymphoid structures (TLSs). TLSs are highly organized aggregates of lymphoid cells that develop in inflamed non-lymphoid tissues and play a critical role in local antigen presentation, T and B cell activation, and anti-tumor immunity. Previous studies have demonstrated that BTK inhibition promotes conventional dendritic cell (cDC) activation in solid tumors, which is essential for initiating adaptive immune responses [Munn, *Immunity*, 2021]. We first characterized TLS formation in GSC005 tumors treated with oHSV, with or without BTK inhibition. TLSs are defined by dense clusters of CD45⁺ immune cells, including B220⁺ B cells and CD3⁺ T cells^24^. Immunofluorescence analysis of tumor sections revealed that treatment with either ibrutinib or oHSV alone did not substantially alter the abundance of B220⁺ B cells compared to mock-treated controls. In contrast, combination treatment with ibrutinib and oHSV induced a shift in TLS composition, characterized by reduced B220⁺ B cell intensity and increased CD3⁺ T cell accumulation, indicating the formation of T cell–rich TLSs (**Fig. 6a–b**). Further staining for MAdCAM-1, a key marker for tumor-associated high endothelial venules (HEVs) for lymphocytes trafficking and TLS formation^25^, revealed elevated expression in tumors receiving combination therapy compared to single-agent treatments (**Fig. 6c**). Mechanistically, ibrutinib co-treatment upregulated the expression of lymphoid chemokines and TLS-associated molecules, including Ltb, Tnfsf14, Ccl19, and Ccl21, which are known to drive TLS neogenesis^25^ and recruitment of immune effector cells (**Fig. 6d**). These findings indicate that oHSV combined with BTK inhibition enhances TLS formation within the GBM microenvironment, creating a supportive niche for T cell–mediated anti-tumor responses.

**Fig. 6.**
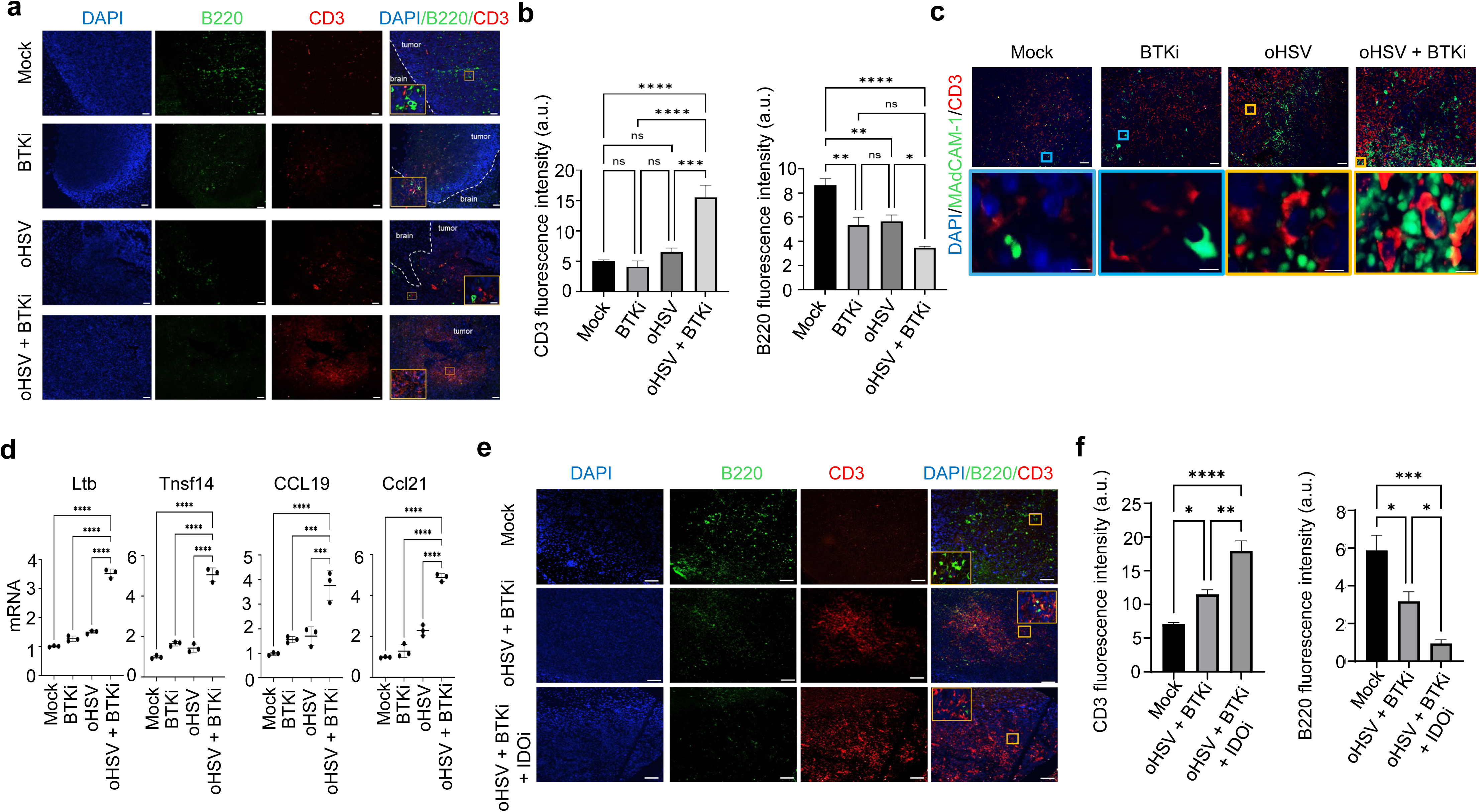
BTK and IDO inhibition promote tertiary lymphoid structure (TLS) formation in murine GSC005 GBM. (**a, c**) Representative immunofluorescence images (**a**) and quantification (**b**) of TLS formation in tumors treated with oHSV, ibrutinib, or their combination, showing increased TLS density with BTK inhibition. n=5, * *p*<0.05, ** *p*<0.01, *** *p*<0.005, **** *p*<0.0001. ns=no significance. (**c**) Representative immunofluorescence images of MAdCAM-1 in tumors treated with oHSV, ibrutinib, or their combination, showing increased TLS density with BTK inhibition. (**d**) qRT-PCR analysis of TLS-associated gene expression in GSC005 tumors following treatment with oHSV alone or in combination with ibrutinib and/or indoximod, demonstrating upregulation of TLS-related signatures. (**d, e**) Immunofluorescence staining (**d**) and quantification (**e**) of TLS in tumors treated with the addition of IDO inhibition, revealing enhanced TLS maturation and organization compared to dual therapy. n=5, * *p*<0.05, ** *p*<0.01, *** *p*<0.005, **** *p*<0.0001.

We further examined whether addition of indoximod could potentiate TLS formation during dual therapy with oHSV and ibrutinib. Triple treatment increased CD3⁺ T cell infiltration while further reducing B220⁺ B cell intensity relative to the dual therapy group (**Fig. 6e–f**), suggesting enhanced T cell–dominated immune structures and improved antigen presentation. Collectively, these results indicate that IDO inhibition reinforces the immunostimulatory effects of oHSV plus BTK inhibition by promoting TLS development, improving local antigen presentation, and fostering a microenvironment conducive to robust anti-tumor T cell responses in GBM.

### Addition of IDO inhibitor to oHSV plus Ibrutinib further remodels the tumor Immune microenvironment

To determine whether the enhanced anti-tumor efficacy of adding indoximod to oHSV plus ibrutinib is associated with remodeling of the tumor immune microenvironment, we performed single-cell RNA sequencing (scRNA-seq) on tumors following treatment. Analysis revealed that the oHSV plus ibrutinib regimen increased overall T cell infiltration within the tumor (**Fig. 7a–b**). Importantly, the addition of indoximod further reshaped T cell phenotypes, including an increase in cytotoxic T lymphocytes (CTLs, Cd8a+Gzmk+Tcf7+) (**Fig. 7b**). Notably, a distinct population of NKT cells (Klra7+Cd7+Tcf7+) emerged in the tumor-infiltrating lymphocytes (TILs) when indoximod was added to the oHSV/ibrutinib regimen (**Fig. 7c**), suggesting that IDO inhibition promotes recruitment or expansion of specialized effector subsets.

**Fig. 7.**
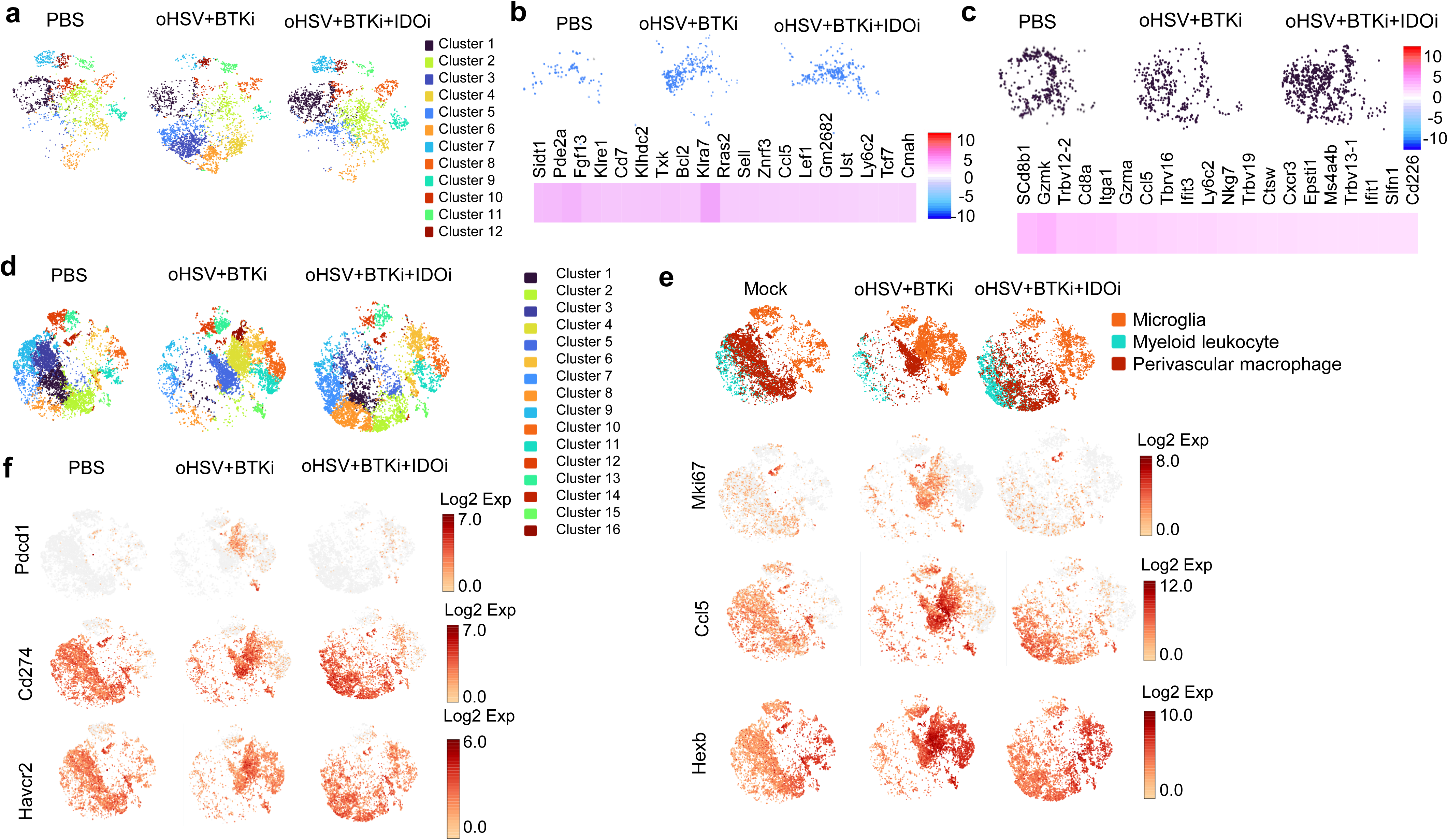
IDO inhibition further reshapes the immune landscape during combined oHSV and BTK inhibition. (**a**) tSNE visualization of T cell populations in murine GSC005 tumors treated with PBS, oHSV plus BTK inhibitor (BTKi), or oHSV + BTKi + IDO inhibitor (IDOi), illustrating distinct immune phenotypic clustering. (**b**) Quantification of effector T cell subsets, showing increased activation and expansion with combination therapies. (**c**) Analysis of natural killer T (NKT) cell populations demonstrating modulation by IDO inhibition. (**d-f**) Identification of mono-cyte-macrophage subclusters by scRNA-seq, revealing significant immune remodeling of inflammation (Mki-67, Ccl5, Hexb), immunosuppression (Pdcd1, Cd274 and Havcr2), and polarization changes following addition of IDO inhibition.

Given that myeloid cells represent the dominant immune population in brain tumors, we next examined monocyte–macrophage subsets, which constitute the major myeloid compartment within the tumor microenvironment. Analysis of these clusters demonstrated that the oHSV/ibrutinib combination reshaped monocyte–macrophage subpopulations, and the addition of indoximod further altered their phenotypic composition (**Fig. 7d**). Sub-clustering of the monocyte–macrophage compartment identified three major populations: microglia, myeloid leukocytes, and perivascular macrophages (pvMacs) (**Fig. 7e**). Both pvMacs and microglia showed increased expression of Ki67, CCL5, and Hexb following oHSV and BTKi treatment, indicating enhanced inflammatory and neurotoxic activity, which was attenuated by the addition of IDO inhibition (**Fig. 7e**). Overall, GBM-associated monocyte–macrophages exhibited an immunosuppressive phenotype marked by PD-1, PD-L1, and TIM-3 expression. oHSV and BTKi further changed these inhibitory markers, particularly in pvMacs and microglia, whereas IDO inhibition partially reversed this effect and reshaped the myeloid immunosuppressive profile (**Fig. 7f**). These changes included shifts toward subpopulations associated with antigen presentation and reduced immunosuppressive activity, consistent with enhanced anti-tumor immunity.

Collectively, these findings indicate that IDO inhibition amplifies the immunomodulatory effects of BTK inhibition during intratumoral oHSV therapy in GBM. By promoting CTL and NKT cell infiltration and further reprogramming monocyte–macrophage phenotypes, indoximod contributes to a more pro-inflammatory, thereby enhancing the overall efficacy of the combinatorial treatment.

## Discussion

Bruton’s tyrosine kinase (BTK) plays a central role in B cell development, and its role in B cell malignancies has been extensively studied. The BTK inhibitor ibrutinib is FDA-approved for multiple B cell malignancies, including chronic lymphocytic leukemia and mantle cell lymphoma. Beyond B cells, BTK is expressed in myeloid cells, where it regulates proliferation, survival, antigen presentation, and immune functions. Glioblastoma (GBM) is a highly aggressive tumor characterized by rapid proliferation, invasion, and resistance to therapy^26–28^. Myeloid cells—including macrophages and microglia—are the predominant immune populations in GBM and contribute to the immunosuppressive tumor microenvironment (TME). In this study, we systematically interrogated the role of myeloid BTK in modulating GBM response to oHSV therapy. Our findings indicate that BTK inhibition enhances oHSV anti-tumor efficacy through both immune-independent mechanisms, by directly enhancing tumor lysis, and immune-mediated mechanisms, by reshaping the TME.

Previous studies reported that ibrutinib suppresses glioblastoma stem cell (GSC) stemness via inhibition of BMX–STAT3 signaling^10^. Consistent with prior work, we observed that ibrutinib alone had modest effects on GBM neurosphere proliferation but did not significantly inhibit oHSV infection or replication. Importantly, the combination of ibrutinib and oHSV enhanced tumor cell lysis *in vitro*, highlighting the capacity of BTK inhibition to potentiate viral oncolysis without impairing viral propagation. This demonstrates that BTK activity modulates tumor susceptibility to oHSV, supporting the rationale for combination therapy in GBM.

GBM-infiltrating macrophages and microglia were identified as the primary BTK-expressing cells in both human and murine GBM models. Intra-tumoral oHSV injection increased myeloid infiltration and BTK activation, which promoted viral clearance via TNF-α–dependent mechanisms and enhanced residual tumor cell invasion. Addition of ibrutinib reduced BTK activity in these myeloid populations, decreasing oHSV clearance while simultaneously inhibiting tumor invasiveness. These findings highlight BTK’s dual role in antiviral defense and tumor-promoting functions within the GBM TME.

BTK is also expressed in tumor-infiltrating dendritic cells, where it modulates antigen presentation. Previous studies have shown that BTK regulates immunosuppressive IDO activity in conventional dendritic cells, and BTK inhibition enhances dendritic cell activation and antigen-specific T cell responses^11^. In our study, ibrutinib increased CD8⁺ T cell infiltration in GBM. When combined with the IDO inhibitor indoximod, oHSV-induced anti-tumor immune responses were further amplified. This included enhanced TLS formation, increased CTLs and NKT cells, and reprogramming of tumor-infiltrating myeloid populations. These results support a model in which BTK inhibition alleviates myeloid-mediated immunosuppression, allowing oncolytic viruses to prime robust adaptive immunity.

IDO is overexpressed in multiple tumor types, including GBM, where it suppresses T cell activity and antigen presentation^11,29–31^. In our previous work, oHSV therapy upregulated IDO in tumor-infiltrating dendritic cells, creating feedback inhibition of antigen presentation. Addition of indoximod reversed this suppression and further enhanced anti-tumor immune responses^23^. Importantly, BTK signaling also regulates IDO via the BTK–IDO–mTOR axis, linking BTK activity to immunosuppressive pathways in antigen- presenting cells. Combining ibrutinib with indoximod and oHSV maximized dendritic cell activation, increased CD8⁺ T cell infiltration, and enhanced tumor cell apoptosis.

TLSs are organized lymphoid aggregates that function as local antigen presentation sites, complementing tumor-draining lymph nodes. TLS facilitates T cell priming and are associated with improved responses to immunotherapy. In our study, combination oHSV and BTK inhibition increased TLS formation within tumors, characterized by CD3⁺ T cell–rich clusters and enhanced expression of HVEM, Ltb, Tnfsf14, Ccl19, and Ccl21. The addition of indoximod further augmented TLS maturation, increased CD3⁺ T cell density, and decreased B220⁺ intensity, reflecting enhanced antigen presentation and adaptive immune activation. These findings suggest that combinatorial modulation of BTK and IDO signaling reshapes the GBM TME to favor robust anti-tumor immunity.

Effective GBM therapy requires overcoming the blood–brain barrier (BBB) and blood–tumor barrier (BTB). Ibrutinib exhibits high BBB permeability due to its low molecular weight and hydrophobicity and can transiently disrupt BTB integrity, facilitating drug delivery and improving survival in glioma models^32,33^. These properties, combined with its immune-modulatory and antiviral-enhancing effects, make ibrutinib an ideal candidate for combinatorial GBM therapy.

Oncolytic viruses are capable of self-replication and tumor-selective lysis but are limited by antiviral responses and the immunosuppressive TME^34^. Combining oHSV with BTK and IDO inhibitors provides a multifaceted approach: (i) enhancing viral oncolysis, (ii) inhibiting immunosuppressive myeloid activity, (iii) promoting TLS formation, and (iv) stimulating CD8⁺ T cell and NKT responses. These synergistic effects reflect a broader principle that combining oncolytic virotherapy with immune and pharmacologic modulators can overcome resistance mechanisms and improve therapeutic efficacy^35–38^. Preclinical models demonstrate that these combinatorial strategies significantly prolong survival, suggesting translational potential for clinical GBM therapy.

In conclusion, BTK regulates both tumor-intrinsic and immune-mediated mechanisms that limit oHSV efficacy in GBM. Targeting BTK with ibrutinib, particularly in combination with IDO inhibition, remodels the immune TME, enhances TLS formation, increases cytotoxic immune cell infiltration, and potentiates viral oncolysis. This study provides strong preclinical evidence supporting the rational design of combinatorial BTK- and IDO-targeted therapies to overcome GBM immunosuppression and improve clinical outcomes.

## Acknowledgement

This work was supported by grants from the National Institute of Health (NIH) (R21NS130429 to BH), (NIH) (R61NS128191 to BH and BK), Alex Lemonade Stand Foundation Reach Grant (ALEX23-27891 to BH), and Paceline Foundation (MCG8451Tto BH). GKF is supported by Rally Foundation for Childhood Cancer Research, CureSearch for Children’s Cancer, V Foundation, Hyundai Hope on Wheels, and Kids Join the Fight.

## Author contributions

BH. designed experiments. KK, DZ, KM, GKF, DL, MW, NFM, RK, TSJ, and BH performed the experiments and analyzed the data. HD performed bulk mRNA-seq and scRNA-seq analysis. BH, BK, and DHM wrote and revised the manuscript. All authors edited and reviewed the manuscript.

---

**sFig. 1.**
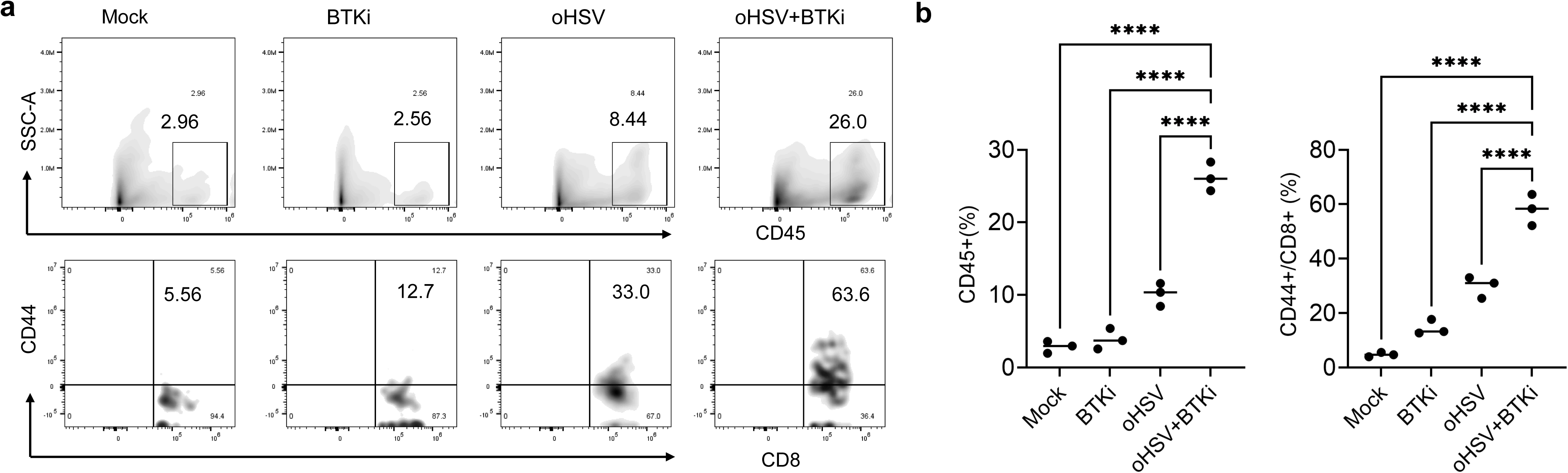
CD8 T cells activation in GSC005 tumor during combined oHSV and BTK inhibition. (**a**) Flow cytometry analysis and quantification (**b**) of CD8 T cell activation in murine 005 tumors treated with PBS, BTK inhibitor (BTKi), oHSV, or oHSV plus BTKi on day 7 after treatment. n=3, **** *p*<0.0001.

